# Wings of Change: aPKC/FoxP-dependent plasticity in steering motor neurons underlies operant selflearning in *Drosophila*

**DOI:** 10.1101/2022.12.16.520755

**Authors:** Andreas Ehweiner, Carsten Duch, Björn Brembs

## Abstract

**Background:** Motor learning is central to human existence, such as learning to speak or walk, sports moves, or rehabilitation after injury. Evidence suggests that all forms of motor learning share an evolutionarily conserved molecular plasticity pathway. Here, we present novel insights into the neural processes underlying operant self-learning, a form of motor learning in the fruit fly *Drosophila*.

**Methods:** We operantly trained wild type and transgenic *Drosophila* fruit flies, tethered at the torque meter, in a motor learning task that required them to initiate and maintain turning maneuvers around their vertical body axis (yaw torque). We combined this behavioral experiment with transgenic peptide expression, CRISPR/Cas9-mediated, spatio-temporally controlled gene knock-out and confocal microscopy.

**Results:** We find that expression of atypical protein kinase C (aPKC) in direct wing steering motoneurons co-expressing the transcription factor *FoxP* is necessary for this type of motor learning and that aPKC likely acts via non-canonical pathways. We also found that it takes more than a week for CRISPR/Cas9-mediated knockout of *FoxP* in adult animals to impair motor learning, suggesting that adult *FoxP* expression is required for operant self-learning.

**Conclusions:** Our experiments suggest that, for operant self-learning, a type of motor learning in *Drosophila*, co-expression of atypical protein kinase C (aPKC) and the transcription factor *FoxP* is necessary in direct wing steering motoneurons. Some of these neurons control the wing beat amplitude when generating optomotor responses, and we have discovered modulation of optomotor behavior after operant self-learning. We also discovered that aPKC likely acts via non-canonical pathways and that *FoxP* expression is also required in adult flies.

## 1. Introduction

Motor learning is an essential component of human behavior and ubiquitous throughout the animal kingdom. The process of learning a motor skill can be influenced by a number of factors, such as the amount of training, type of feedback, or the presence/absence of environmental cues. Regaining lost motor functions after brain or spinal cord injury is considered a crucial component of rehabilitation. Human language is acquired by a form of motor learning, and motor learning also appears to be a key invention that allowed the newly evolved animals in the Cambrian to become ambulatory (1–3). Vocal learning, such as learning to speak, is a form of motor learning (4) that involves the Forkhead Box transcription factor family P (FoxP) in vertebrates (5–15). Other, nonvocal forms of motor learning also involve FoxP genes in vertebrates (16–19) and invertebrates (20). Vocal learning also shares the involvement of protein kinase C (PKC) with other forms of motor learning (21–25), raising the possibility of a conserved motor learning pathway extending beyond these two components.

While motor learning shares various features with other forms of learning, such as operant learning producing habits or skill-learning, it is debated how many common biological mechanisms these different concepts share (26–29). Understanding motor learning in a numerically smaller nervous system in a genetically tractable organism where one can not only study the motor learning process itself, but also its interactions with other forms of learning (30), may help inform these debates.

Here we provide further evidence about the specific manner in which FoxP and PKC are involved in a form of motor learning in the fruit fly *Drosophila*, operant self-learning at the torque meter. In this experiment, motor learning dissociates from other forms of learning such that genes involved in motor learning are not involved in other forms of learning and *vice versa* (31–33). At the torque meter, a fly is tethered between head and thorax such that it can move all other appendages. When beating its wings, the fly generates forces, some of which can be measured by the torque meter. Specifically, the torque meter measures torque around the vertical body axis, yaw torque (34). Even in the absence of any guiding cues, flies can learn to associate one torque domain (e.g., roughly corresponding to left or right, respectively, turning maneuvers) with a punishing heat beam (35). This experiment not only conceptually mimics other motor learning paradigms in that feedback is made immediately contingent on specific motor actions, but also via its dependence on *FoxP* and PKC genes (20,21). This form of motor learning has been termed operant self-learning to distinguish it from other forms of operant learning and to denote that the subject is learning about its own behavior, as opposed to some stimulus associated with the behavior (32).

For operant self-learning in *Drosophila*, it is not known in which neurons the *FoxP* gene is required and which PKC gene is involved. It is also unknown which pathway is engaged by PKC and whether FoxP expression is also required acutely in adult flies for this form of motor learning. In this work, we addressed all three research questions.

## 2. Methods

### 2.1. Strains and fly rearing

If not stated otherwise, flies (table 1) were raised on standard cornmeal/molasses medium (36) at 25°C and 60% humidity under a 12-hour light/dark cycle. For experiments requiring the expression of temperature-sensitive Gal80, animals were raised at 18°C. To set up crosses for behavioral experiments, 20 females were placed together with five to eight males and were allowed to lay eggs for 24 h. They were flipped daily into fresh vials, to ensure appropriate larval density. In 30 years of research on learning and memory in tasks like ours, no difference was ever observed between male and female flies in terms of learning ability. Whenever genetically appropriate, we used female flies for practical reasons. The flies were prepared the day before the experiment, allowing them time to recover. Female flies (24 to 48 h old) were briefly immobilized under cold anesthesia. A thin triangular copper hook (0,05 mm diameter) was glued (3m Espe Sinfony, 3M Deutschland GmbH) between head and thorax, fixing both body parts to each other (37). Each animal was kept individually in a small moist chamber with a few grains of sugar. For *tub-Gal80^ts^* expression, animals were raised at 18°C and incubated at 30°C for two days. Experiments were always conducted at room temperature. For experiments using the gene-switch system, newly hatched flies were placed on *Drosophila* instant medium (351.204, Schlüter Biologie, Eutin-Neudorf, Germany) containing the steroid hormone RU486 (200 μg/ml, CAS No.: 84371-65-3, Sigma-Aldrich, St. Louis, MO) for two days.

**Table 1:** Fly strains used in this work.

| Genotype | use | Bloomington | Flybase |
| --- | --- | --- | --- |
| ;;nSyb-GS | pan-neuronal driver | 80699 | FBti0201287 |
| ;;GMR65A06-GAL4 | protocerebral bridge driver | 39330 | FBti0137511 |
| ;;GMR20H05-GAL4 | central complex driver | 47896 | FBti0133817 |
| ;;ato-Gal4 | driver for dorsal cluster neurons |  |  |
| ELAV-Gal4;; | pan-neuronal driver |  |  |
| ELAV-Gal4;Tub-Gal80ts;; | temperature-sensitive pan-neuronal driver |  |  |
| ;;FoxP-iB-Gal4/TM3 | driver for FoxP-iB neurons |  |  |
| ;;FoxP-LexA | driver for all FoxP neurons |  |  |
| ;;UAS-g-aPKC | effector | 85862 | FBti0210993 |
| ;;UAS-g-BAZ | effector | 84234 | FBti0207133 |
| ;;UAS-g-PKC53e | effector | 76612 | FBti0194968 |
| ;UAS-g-KIBRA | effector |  |  |
| ;;UAS-Cas9; | effector |  |  |
| ;;UAS-PKCi | effector | 4589 | FBti0010565 |
| ;;UAS-t:gRNA(4xFoxP) | effector |  |  |
| ;UAS-Cas9;; | effector |  |  |
| C380-Gal4;; | MN driver | 80580 | FBti0016294 |
| ;;D42-Gal4 | MN driver | 8816 | FBti0002759 |
| y[1] w[*]; Mi{Trojan GAL4.un} aPKC[MI10848-TG4.un]/SM6a | driver for aPKC cells | 77814 | FBti0196316 |
| LexAop-mD8-RFP-UAS-mCD8-GFP;TM3/TM6b | effector |  |  |
| Wild type Berlin | wild type strain |  |  |

The UAS-g-KIBRA line was generated in the laboratory of Michael Krahn, Universität Münster. They cloned the following guide RNAs from KIBRA Intron-2: GTACTTAC-GACTGCTTCGAC and KIBRA Intron-4: GGGCAC-CGTGCAGATCAGCA in pCDF6 and inserted them in attP40.

### 2.2. Behavioral setups

Two different torque meters and setups had to be used for technical reasons, not by choice. Both torque meters were described previously: The ‘Tang’ meter (38) and the ‘Götz’ meter (34)(RRID:SCR_017276), named after the authors. With flies attached to the torque meter via a clamp, all devices measure the rotational force (torque) around the animal’s vertical body axis. While the Götz meter is older than the Tang meter, it is technically more advanced because of its rotational compensation, and was included in a more modern version of the setup that was used in the later experiments, after the first data were collected using the Tang setup. Even later, the wild type experiments shown in Figure 4 were conducted with a third torque meter, still in the prototype phase, which combines the laser-based measurement of the Tang meter with induction-based compensation of the Götz meter. Documentation of this setup is in preparation, but the operating principles remain the same as for the two referenced devices. In all setups, the animal is surrounded by a cylindrical panorama (arena; diameter 58 mm Tang, 90 mm other setups), homogeneously illuminated from behind by either a projector (new setups: DLPLCR4500EVM, Texas Instruments) or a halogen lamp (Tang: OSRAM 100W/12V), such that stationary flight in a controlled environment was achieved. An infrared laser (Stocker Yale Lasiris SNF series; 825 nm, 150 mW) was used as punishment in all setups. The laser was pointed from above onto the animal’s head, pulsed (approximately 200 ms pulse width ∼4 Hz) and the intensity adjusted empirically for maximal heat avoidance and learning. The experiment is fully computer controlled, using custom software (Tang: Lab-View, National Instruments, RRID:SCR_014325. New setups: DOI: 10.5281/zenodo.7102195).

### 2.3. Design of behavioral experiments

Before each self-learning experiment, the yaw torque range was adjusted using optomotor stimuli for each fly tethered to the torque meter. The optomotor response (OMR) is an innate, orienting behavior evoked by whole-field visual motion and is common to vertebrates and invertebrates. The OMR has algorithmic properties such that the direction of the whole-field coherent motion dictates the direction of the behavioral output (e.g., leftward visual stimuli lead to turning left, and rightward visual stimuli lead to turning right). For instance, when tethered Drosophila are surrounded by vertical black and white grating patterns rotating along the fly’s azimuth (i.e, around the fly’s vertical body axis), the fly will turn (i.e., produce yaw torque) in the direction of perceived motion. Typical OMRs for tethered flies responding to horizontally rotating vertical stripes are depicted in Figure 4A1.

For the ‘Tang’ setup, arena rotation for the optomotor stimulus was operated by hand. The direction of the rotation was reversed after the fly reached its asymptotic optomotor torque. During optomotor presentations before the self-learning experiment, the torque was adjusted to be zero-symmetric. This was to facilitate unbiased torque preferences before training. Torque traces during OM presentations were not stored in experiments using the ‘Tang’ setup. Optomotor stimuli (15 vertical black stripes on a white background taking about 3.5s for a full rotation, i.e., a pattern wavelength of 24° at a pattern frequency of about 4.3 Hz) were presented for a duration of 30 s in each turning direction for flies in the new setups and recorded in the raw data files together with all other data from each experiment. Because of this difference, as experiments recorded with the Tang device did not record optomotor periods, periods in ‘Tang’ experiments are numbered from 1 to 9 (Table 2), while optomotor periods for the new setups were included, such that the periods in the new setups were numbered 1-17 (Table 3). Because we used four optomotor periods each before and after training in the new setup, periods 5 to 13 were the periods in which self-learning was studied in this setup.

**Table 2:** Experimental sequence “Tang Setup”. All periods lasted 120s.

| Period 1 | Period 2 | Period 3 | Period 4 | Period 5 | Period 6 | Period 7 | Period 8 | Period 9 |
| --- | --- | --- | --- | --- | --- | --- | --- | --- |
| Pretest | Pretest | Training | Training | Test | Training | Training | Test | Test |
| No heat | No heat | Heat | Heat | No heat | Heat | Heat | No heat | No heat |

**Table 3:** Experimental sequence for new setups. OM - optomotor. Torque learning periods (5–13) lasted 120s, while OM periods lasted 30s.

| OM before |  |  |  | Experiment |  |  |  |  |  |  |  |  | OM after |  |  |  |
| --- | --- | --- | --- | --- | --- | --- | --- | --- | --- | --- | --- | --- | --- | --- | --- | --- |
| Period 1 | Period 2 | Period 3 | Period 4 | Period 5 | Period 6 | Period 7 | Period 8 | Period 9 | Period 10 | Period 11 | Period 12 | Period 13 | Period 14 | Period 15 | Period 16 | Period 17 |
| OM | OM | OM | OM | Pretest | Pretest | Training | Training | Test | Training | Training | Test | Test | OM | OM | OM | OM |
| No heat | No heat | No heat | No heat | No heat | No heat | Heat | Heat | No heat | Heat | Heat | No heat | No heat | No heat | No heat | No heat | No heat |

The main self-learning experiment then consisted of nine periods of two minutes duration in both setups. The laser was permanently off during the first two periods, so that the fly could freely choose its direction of turning maneuvers without any feedback. In the following two training periods either the left or the right torque domain was associated with the punishing laser, without any hysteresis. The punished torque domain was alternated between experiments. The first two training periods were followed by one test period without punishment. Afterwards, the fly was trained again with the same side punished as before for another two 2-min. periods. Finally, no heat was applied in the final two test periods, allowing the fly to express its spontaneous yaw torque preference. The figures always show the preference in the first test period after the last training period, i.e., period 8 (performance index, PI8) in the Tang setup and period 12 (PI12) in the new setups (Tables 2, 3). When the axis labels in the figures differ with regard to PI8 or PI12, these differences only indicate which setup was used. In all cases, the same first test period after the last training period was used to test for learning, irrespective of setup.

### 2.4. Data handling and statistical analysis of behavioral experiments

#### 2.4.1. Data selection

To ensure proper punishment by the laser, each fly was exposed to the laser after the experiment, to ensure it was adjusted correctly. If the fly survived the laser for 15 s or longer, the data were excluded from analysis. Data that did not show any or shifted OMRs, indicating either an unhealthy fly or an error with the measuring device, were also excluded. Data were also excluded if the fly had not ex-perienced the laser at least once during training. Finally, flies with poor flight performance (constant stopping of flight) were also excluded from analysis. While these data were excluded from analysis, all complete traces are nevertheless included in the published data sets, such that the inclusion criteria can be independently tested.

#### 2.4.2. Data availability and analysis

The preference of a fly for right or left torque domain was quantified as the performance index *PI* = (*ta* − *tb*)∕(*ta* + *tb*). During training periods, *tb* indicates the time the fly is exposed to the heat and *ta* the time without heat. During tests, *ta* and *tb* refer to the times when the fly chose the formerly (or subsequently) unpunished or punished situation, respectively. Thus, a PI of 1 indicates the fly spent the entire period in the situation not associated with heat, whereas a PI of −1 indicates that the fly spent the entire period in the situation associated with heat. Accordingly, a PI of zero indicates that the fly distributed the time evenly between heated and nonheated situations and a PI of 0.5 indicates that 90 of the 120 s in that period were spent in the unpunished situation.

Analogously, optomotor (Table 3, OM) behavior was quantified by computing an OM asymmetry index from the torque traces. Straight lines or double sigmoidal models were fitted to individual torque traces from OM periods, depending on the detected slope of the OMR. Each fit was generated separately for each turning direction. From the fitted lines/models, optomotor magnitude was derived as either the intercept (lines) or the asymptote (double sigmoidal model) for each turning direction. The magnitude of left-turning torque was subtracted from the magnitude of right-turning torque and divided by the sum of the two values. This optomotor asymmetry index becomes -1 for OMRs where clockwise (‘right-turning’) stimuli elicit no or left-turning torque (while counter-clockwise stimuli elicit left-turning torque). It becomes 1 for OMRs where counter-clockwise (‘left-turning’) stimuli elicit no or right-turning torque (while clockwise stimuli elicit right-turning torque). The OM asymmetry index becomes zero if the absolute magnitudes of OMRs in both directions are equal. In brief, a positive optomotor asymmetry index indicates shifts away from symmetrical torque towards right-turning torque and a negative index indicates shifts towards left-turning torque.

All behavioral data were analyzed using R (R Project for Statistical Computing) (RRID:SCR_001905). The collection of R-scripts evaluating the time series data can be found at DOI 10.5281/zenodo.10041052 (39). The data model pertaining to the XML raw data files and the YAML data set files can be found at 10.5281/zenodo.7101734 (39). In brief, the XML data files contain both the meta-data for each single fly experiment as well as the time-series data covering the entire experiment. The single YAML file per dataset contains the experimental design, such as which data files belong to which experimental group, the type of statistics to be performed, significance levels used, experimenter comments and data inclusion/exclusion. The main R-Script reads the YAML dataset files and performs the appropriate computations for quality control, analysis and statistics.

Quality control is performed on each single-fly XML file and included in the published datasets. Each single-fly experiment XML file is thus accompanied by a single-fly HTML quality control report sheet containing plots of the raw time series data, as well as a number of evaluations necessary to assess the proper execution of the experiment and the quality of the resulting data.

Data analysis and statistics for each dataset are reported in an HTML dataset evaluation sheet. Thus, a complete dataset consists of one XML raw data file and one HTML quality control sheet for each fly, plus a single YAML dataset file and one HTML dataset evaluation sheet.

The datasets were published using a custom Python script (DOI: 10.5281/zenodo.7101741)(39) that synchronizes the collected data on the local computer with the University of Regensburg publication server. The persistent identifiers for each dataset are listed in the figure legends.

#### 2.4.3. Statistics

Motivated by ongoing efforts to improve statistical inference in science (e.g., (40–54), we chose to statistically evaluate PIs in two complementary ways, using both frequentist and Bayesian statistics (55–61). Following previous studies (30,31,62–65), individual PIs of the first test after the last training period were tested against zero to evaluate the ability of the manipulated flies to show a preference towards the unpunished torque domain. The rationale behind estimating a group of flies as either showing learning or not is to trade-off statistical power with a more nuanced measure of learning performance: comparing between experimental groups may yield more nuance, but also requires impractically large sample sizes for adequate (>80 %) statistical power. To further reduce the chance of statistical error, we used both Wilcoxon tests in a frequentist scenario and computed the equivalent Bayes Factors for a Bayesian version. We set the alpha value for the Wilcoxon test to 0.5 % as suggested by (46), such that p-values below 0.005 and Bayes factors above 5 for the same group of flies would be considered compelling evidence that the flies were able to learn. Conversely, Bayes Factors below one together with p-values higher than 0.05 were considered evidence that the flies were not able to show self-learning. Finally, groups where the two statistics were in conflict or intermediate, were considered inconclusive. Thus, both the Bayesian and the frequentist criteria had to be met in order to claim that a genetic manipulation interfered with operant self-learning. These criteria were chosen to quantitatively distinguish between effective and ineffective manipulations without neglecting the uncertainty associated with all experimentation. At the same time, our evaluations were chosen to specifically identify large contributions to the learning processes that can be identified with sufficient statistical power (i.e., large effect sizes). All statistical results are published with the raw data and the code used to compute them is openly available.

### 2.5. Gene expression analysis in flight steering motor neuron terminals

#### 2.5.1. Dissection

The axon terminals on flight steering muscles were tested immunocytochemically for the expression of *FoxP* and aPKC in flies expressing LexAop RFP under the control of FoxP-Lex and UAS td-GFP under the control of aPKC-Trojan GAL4 (LexAop RFP, UAS td-GFP / aPKC-Trojan-GAL4; FoxP-LexA / +). In addition, synapses at neuromuscular junctions were labeled with the active zone marker *bruchpilot* (*brp*, (66)). Animals were dissected in normal saline along the dorsal midline and bent open with minute pins inserted through the dorsalmost edge of the dorsal longitudinal flight muscles (DLMs). The heart, gut, fat tissue, and other connective tissue were removed to expose the ventral nerve cord (VNC) and the musculature. Next, on both sides the DLMs and dorsoventral flight muscles (DVMs) were carefully removed layer-by-layer to expose the direct flight steering muscles, which are located close to the lateral cuticle of the thorax. Preparations were rinsed 5-10 times in saline to remove debris from the dissection procedure, and specimens were fixed for 1 h in 4 % paraformaldehyde in 0.1 M PBS buffer at room temperature (22°C). After fixation specimen paraformaldehyde was exchanged with 0.1 M PBS buffer.

#### 2.5.2. Immunohistochemistry of motor terminals

Following fixation, specimens were washed 6 × 20 min in 0.1 M PBS buffer at room temperature. Next, preparations were washed 3 × 1 h in 0.1 M PBS-Tx (0.3 %) buffer at room temperature and incubated with primary antibodies (mouse α-brp, 1:500 (Hybridoma bank, NC82)); chicken α-GFP (1:1000, Life Technologies A10262), and rabbit α-mCherry, (1:500, PAS-34974)) in 0.1 M PBS-Tx (0.3 %) buffer at 4° C for 24 to 36 h. Following primary antibody incubation, preparations were washed 6 × 1 h in 0.1 M PBS buffer at room temperature. Next animals were incubated in secondary antibodies: donkey α-mouse Alexa 647 (JacksonImmunoResearch 715-605-150), donkey α-chicken Alexa 488 (Dianova 703-545-155), and donkey α-rabbit Alexa 568 (Invitrogen A10042) in 0.1 M PBS-Tx (0.15%) buffer at 4° C for 24 to 36 h. Following secondary antibody incubation, preparations were washed 3 × 1 h in 0.1 M PBS buffer at room temperature, dehydrated in an ascending ethanol series (50, 70, 90, and 2 × 100 % EtOH, 15 min each), and cleared for 5 min in methyl salicylate. Finally, preparations were mounted in methyl salicylate in between two coverslips that were glued onto both sides of a round hole (10 mm diameter) drilled into custom-made metal slides of 188 µm thickness (67). Briefly, one cover slip was fixated with superglue underneath the hole, the space was filled with methyl salicylate, the preparation transferred into the mounting media, and another coverslip was carefully placed on top of the hole, so that no air remained in the methyl salicylate filled hole. The top coverslip was fixed to the metal slide using transparent nail polish (DM Markt, Mainz, Germany). After 20 min to let the nail polish dry the preparation was transferred to the microscope. In total 11 animals were subjected to immunocytochemistry on direct flight muscles. In 9 of these 11 preparations, we were able to identify only subsets of the direct wing steering muscles under investigation. These subsets yielded identical results with respect to aPKC and FoxP expression as the two preparations with complete sets of direct muscles as shown in Figure 3.

#### 2.5.3. Confocal laser scanning microscopy

Preparations were scanned using a Leica (Leica Microsystems, Germany) SP8 confocal laser scanning microscope (CLSM) under either a 20x oil (NA = 0.75) or a 40x oil (NA = 1.3) lens at an image format of 1024 × 1024 pixels. Z-step size was 2 µm for the 20x lens (resulting in voxel dimensions of 0.57 × 0.57 × 2 µm) and 1 µm for the 40x lens (resulting in voxel dimensions of 0.28 × 0.28 × 1 µm). Alexa 488 was excited with an argon laser at 488 nm and detected with a photomultiplier between 495 and 530 nm wavelength. Alexa 488 was excited with a solid state laser at 561 nm and detected with a photomultiplier between 570 and 610 nm wavelength. Alexa 647 was excited with a red helium neon laser at 633 nm and detected with a photomultiplier between 640 and 670 nm wavelength. All image stacks were stored as .lei files and further analyzed using Las X software (Leica Microsystems, Germany). Selected fields of view were used for maximum intensity projection views, which were exported as 24 bit three color tiff images and further processed with <u>Corel Draw 11</u>.

#### 2.5.4. Data availability

Confocal image stacks can be found at: 10.5281/zenodo.10606166 (68).

## 3. Results

### 3.1. Self-learning requires a*PKC* in *FoxP* motor neurons

Colomb and Brembs (2016) discovered that blocking all protein kinase C (PKC) isoforms in neurons using the inhibitory peptide PKCi abolished operant self-learning. We replicated their results by pan-neuronal expression of PKCi. In one group, PKCi was expressed both during development and in the adult flies. In the other group, PKCi expression was restricted to adulthood only, by using the temperature-sensitive Gal4 inhibitor *tub-Gal80^ts^* (see Methods for details). Temporally unrestricted pan-neural expression did not impair self-learning, whereas expression restricted to neurons in adult flies abolished self-learning (Figure 1A). This result may seem surprising, but compensation for experimental manipulations of PKC activity through development has been reported numerous times in the literature (69–73) and our data reproduced our previously published, identical experiments with PKCi (21), demonstrating the PKCi construct is still performing as expected. The same publication also reported that PKCi expression in MNs was sufficient to impair operant self learning. The expression of the driver lines used in Colomb and Brembs overlap with the expression of FoxP in the MNs of the ventral nerve cord (74), so we tested if driving PKCi in neurons expressing the isoform B of FoxP was sufficient to impair operant self-learning. Corroborating the notion that MNs are crucial for operant self-learning, expressing PKCi only in cells expressing isoform B of the FoxP gene also abolished self-learning, even without temporal control, suggesting that PKC activity is required in FoxP-positive MNs of the ventral nerve cord.

**Figure 1:**
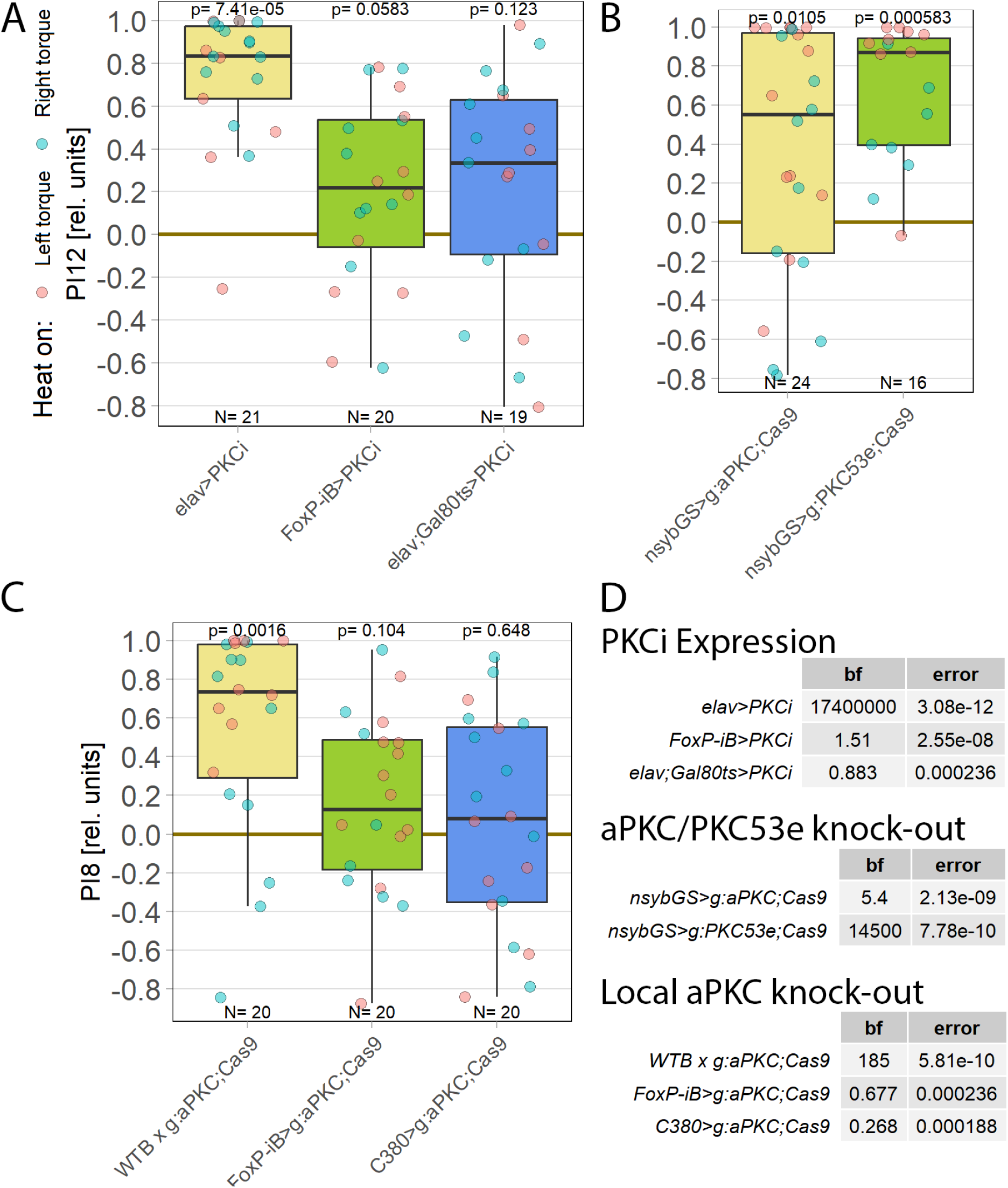
For operant self-learning, *aPKC* is required in motor neurons. Performance indices for the first period after training (PI8/PI12) are plotted. Each colored dot represents a single fly experiment. Red dots denote experiments where the fly was punished on its left turning torque domain, blue dots denote flies that were punished on their right turning domain. Box plots denote medians, quartiles and non-outlier range. Statistical analyses test for differences of PIs against zero. A. Inhibiting all protein kinase C isoforms with the inhibitory peptide PKCi. Constitutive, pan-neuronal expression PKCi (left, yellow), leads to high PIs and a large Bayes Factor, indicating this manipulation left self-learning intact. Expressing PKCi either in FoxP-isoform B positive neurons without temporal control (middle, green), or in all neurons but restricted to adulthood using *tub-Gal80^ts^* (right, blue, see Methods for details) yields low PIs, high p-values and low Bayes factors, indicating self-learning was impaired. Data: 10.5283/epub.52958 (75) B. Pan-neuronal knock-out of two different PKC genes with CRISPR/Cas9 in adulthood (using the GeneSwitch system and feeding RU486 to adult flies, see Methods for details) suggests aPKC is necessary for operant self-learning. Knocking out atypical PKC (yellow, left) yields moderate PIs, p-values and Bayes factors, indicating some effect on operant self-learning, while the high PIs, low p-values and high Bayes factor of the group where PKC53E was knocked out (right, green) indicate their selflearning was intact. Data: 10.5283/epub.52957 (76) C. Knocking out aPKC in motor neurons or *FoxP*-neurons impairs operant self-learning. Expressing the CRISPR/Cas9 components either in *FoxP* isoform B-positive neurons (green, middle) or in motor neurons (blue, right) leads to low PIs, high p-values and low Bayes Factors, indicating their self-learning is strongly impaired. Control flies with only the CRISPR/Cas9 genetic elements but no driver, showed high PIs, a low p-value and a high Bayes Factor, indicating their self-learning was intact. Data: 10.5283/epub.52944 (77) D. Bayesian statistics for the three datasets.

To help determine which PKC may be involved in this mechanism, we screened RNA-Seq databases for PKC genes expressed in MNs. Restricting candidates to those where gRNA lines were available for CRISPR/Cas9-mediated gene knockout yielded only two genes: the atypical PKC (aPKC) and the diacylglycerolactivated PKC53E. Knocking out each gene pan-neuronally in adult flies and testing the manipulated animals for operant self-learning showed excellent learning performance in PKC53E-manipulated flies (Figure 1B), clearly ruling out PKC53E as the gene involved in operant self-learning. These results also confirmed prior experiments with PKC53E mutant flies (21), ruling out any PKC53E involvement. In contrast to the PKC53E manipulated flies, the performance of the aPKC-manipulated flies was inconclusive: their preference scores were somewhat lower, not reaching our criteria for significant learning, but at the same time too high to be confident in the result (Figure 1B). To minimize the possibility of a false-positive result, we replicated the aPKC experiments with different driver lines.

As PKC activity is required in MNs (21), we limited the aPKC knockout to these neurons which abolished self-learning (Figure 1C). To test the hypothesis articulated above that aPKC activity is required in *FoxP* neurons, we also knocked out aPKC in *FoxP* neurons, which also abolished self-learning. (Figure 1C). Using two different driver lines also controls for driver-specific effects and potential expression outside of motor neurons. Both driver lines support operant self-learning in principle (article in preparation, data at DOI: 10.5283/epub.52962), with C380-Gal4 also already in the peer-reviewed literature (21). To our knowledge, C380 and FoxP-iB expression overlaps only in MNs. Thus, three manipulations of aPKC showed effects on self-learning, while two manipulations of PKC53E both failed to show an effect on self-learning.

### 3.2. *aPKC* and *FoxP* are co-expressed in identified flight steering MNs

Thus, the behavioral data presented above support the hypothesis that the plasticity mediating operant self-learning takes place in neurons that co-express both *FoxP* and *aPKC*. Whole-mount confocal microscopy of fly central nervous systems with *aPKC*-Gal4 and *Foxp*-LexA expression suggested that neurons expressing both *aPKC* and *FoxP* exist only in the ventral nerve cord (VNC, Figure 2A, B). We identified co-expressing neurons in all neuromers of the VNC, with the ventral location of the mesothoracic *aPKC/FoxP* neurons suggesting potential wing MNs (Figure 2C)(78). To ascertain the identity of this ventral cluster of co-expressing neurons, we marked MNs with the driver line D42-Gal4 and used FoxP-LexA to stain all *FoxP* neurons. With these labels, we identified a ventral sub-population of putative wing MNs expressing *FoxP*, which matched the location of the *aPKC/FoxP* neurons identified before (Figure 2D). The overlap (or lack thereof) may not be clearly visible in these 2D renderings presented in this text, which is why we made the 3D image stacks available for closer scrutiny (DOI: 10.5281/zenodo.10047941)(79). Both the C380-Gal4 driver used in Figure 1 and the D42-Gal4 driver used here, label not only MNs but also other neurons, in particular cholinergic neurons. However, the lines overlap in MNs in the VNC. This is the main reason why we have both historically and also in this work used both drivers (c380-Gal4 and D42-Gal4) interchangeably. *FoxP* is also expressed in these MNs and knocking out aPKC in these neurons also impaired operant self-learning. None of these lines of evidence is sufficient to conclude that steering MNs are the site of plasticity for operant self-learning. Taken together, they justify testing the hypothesis that wing steering MNs expressing both aPKC and FoxP may be important for operant self-learning. These suggestive results, together with the recently published draft VNC connectome (80), motivated us to analyze the direct wing steering muscles for innervation by MNs with both aPKC and *FoxP* expression, instead of further quantifying the VNC dataset. The expected outcome of this analysis was to obtain a more high-quality and higher resolution picture of aPKC/FoxP expression in direct steering muscle MNs.

**Figure 2:**
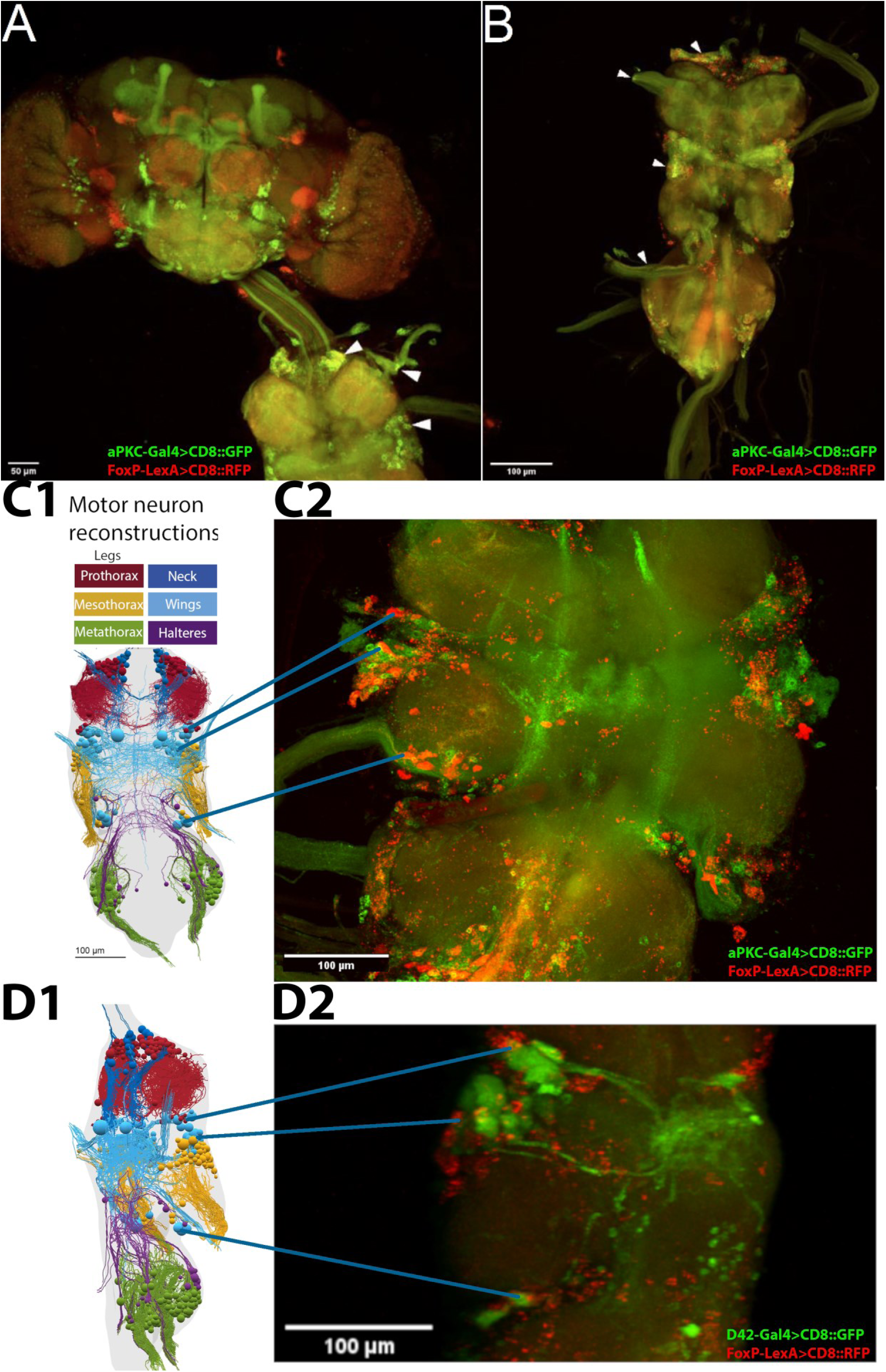
FoxP and aPKC are co-expressed in MNs. Confocal stacks of whole mount preparations of central nervous systems; A-C: green - aPKC-Gal4>CD8::GFP, red - FoxP-LexA>CD8::RFP; D: green - D42-Gal4>CD8::GFP, red - FoxP-LexA>CD8::RFP. Confocal image stacks available at: 10.5281/zenodo.10047941. A. Adult brain (top) with ventral nerve cord (VNC, bottom) attached. No co-expressing cells can be observed in the brain, whereas such neurons(yellow) are readily observable in all neuromers of the VNC (arrowheads). B. VNC with *aPKC/Foxp* co-expression (yellow) both in cell bodies and fiber tracts in nerves (arrowheads). C. C1: Dorsal view of motor neuron reconstruction (modified from (78). C2: Confocal image stack of dorsal view of the mesothoracic neuromer with putative wing MNs expressing both *aPKC* (green) and *FoxP* (red) marked. D. D1: Lateral view of motor neuron reconstruction (modified from (78). D2: Confocal image stack of mesothoracic neuromer with all MNs (green) and FoxP neurons (red) marked. This lateral view supports the hypothesis that the ventral cluster of *aPKC/FoxP* neurons comprises wing MNs.

**Figure 3.**
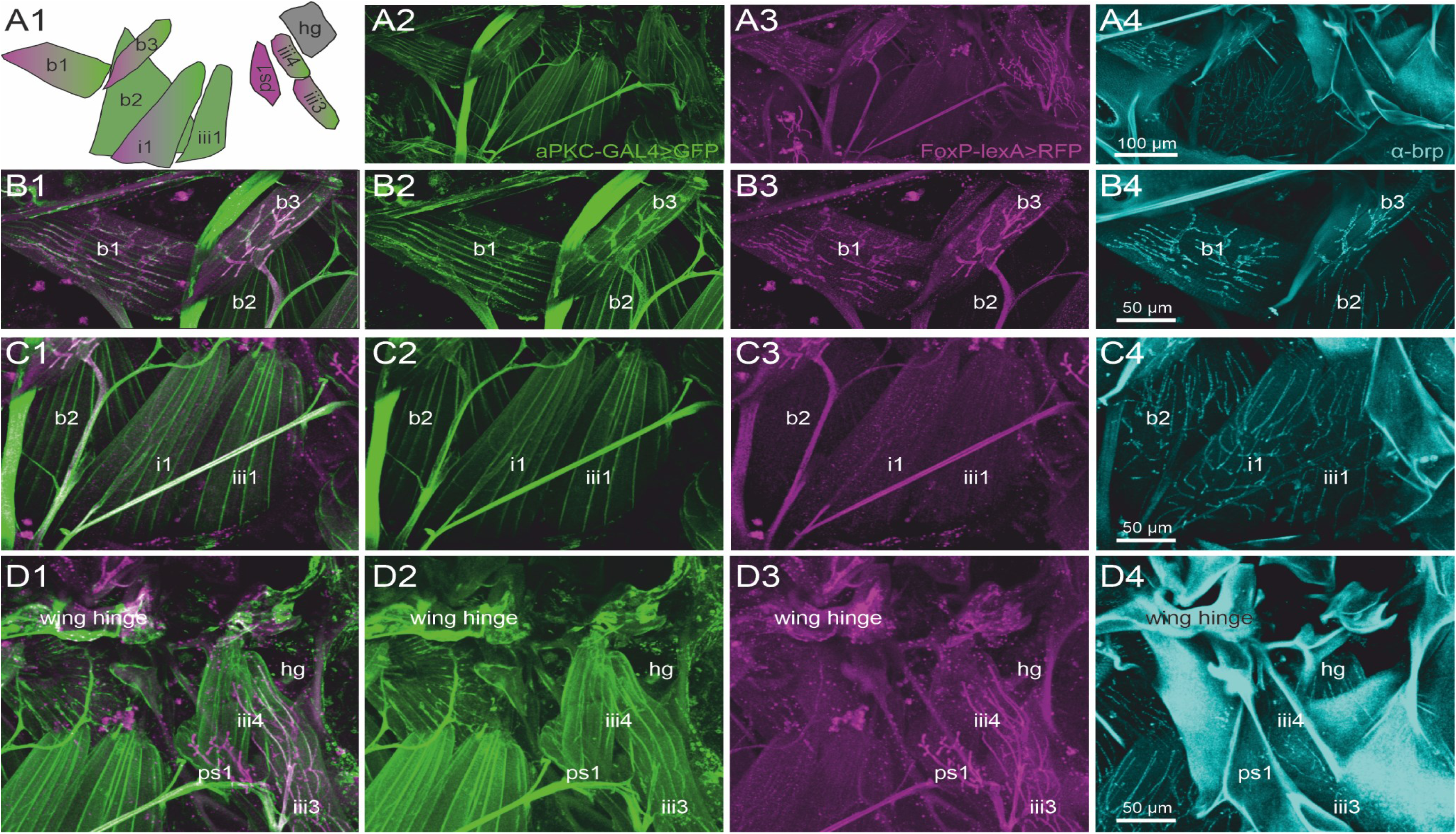
A subset of direct wing muscles is innervated by aPKC/FoxP co-expressing MNs. Representative projection view of MN terminals on the direct flight steering muscles in animals with GFP label in aPKC expressing cells (aPKC-Gal4>CD8::GFP, green), RFP expression in FoxP expressing cells (FoxP-LexA>CD8::RFP, magenta), and immunolabeling for the presynaptic active zone marker *bruchpilot* (*brp*, cyan) reveal which direct flight steering MNs express either aPKC, FoxP, or both, or none of them but only *brp* in presynaptic active zones. **(A1)** depicts the orientation, shape, and abbreviated names of direct flight steering muscles and summarizes which ones are innervated by aPKC expressing MNs (green), by FoxP-expressing MNs (magenta), or by MNs without FoxP and aPKC expression (grey). **(A2)** Projection view of direct flight muscles and their innervation with GFP expression under the control of aPKC-GAL4 (green) at 20x magnification. **(A3)** Same preparation, image stack, and field of view but with RFP expression under the control of FoxP-lexA (magenta). **(A4)** Same preparation, image stack, and field of view with *brp* immunolabel (cyan) in presynaptic active zones of flight steering MNs. **(B1-B4)**. Same preparation but with selective enlargement of the three basalare muscles (b1-b3), with all three labels in **(B1)**, GFP label in aPKC-expressing cells (green, B2), RFP label in FoxP expressing cells (magenta, B3), and Brp label in presynaptic active zones (cyan, B4). Muscles b1 and b3 are innervated by steering MNs with aPKC and FoxP expression, but b2 is devoid of FoxP-expressing innervation. **(C1-C4)** Same preparation but with selective enlargement of second basalare (b2) and the adjacent pterale 1 (i1) and pterale II (iii3) muscles with all three labels **(C1)**, GFP label in aPKC-expressing cells (green, C2), RFP label in FoxP-expressing cells (magenta, C3), and *brp* label in presynaptic active zones (cyan, C4). Only i1 is faintly labeled for terminals with aPKC FoxP expression. **(D1-D4)** Same preparation but with selective enlargement of the pterale II muscles iii3 and iii4, the adjacent pleurosternal muscle (ps1), and the posterior notal wing process muscles (hg) with all three labels **(D1)**, GFP label in aPKC-expressing cells (green, D2), RFP label in FoxP-expressing cells (magenta, D3), and Brp label in presynaptic active zones (cyan, C4). The pterale II muscles iii3 and iii4 are innervated by terminals with aPKC and FoxP expression. Image stacks available at: 10.5281/zenodo.10606166

**Figure 4.**
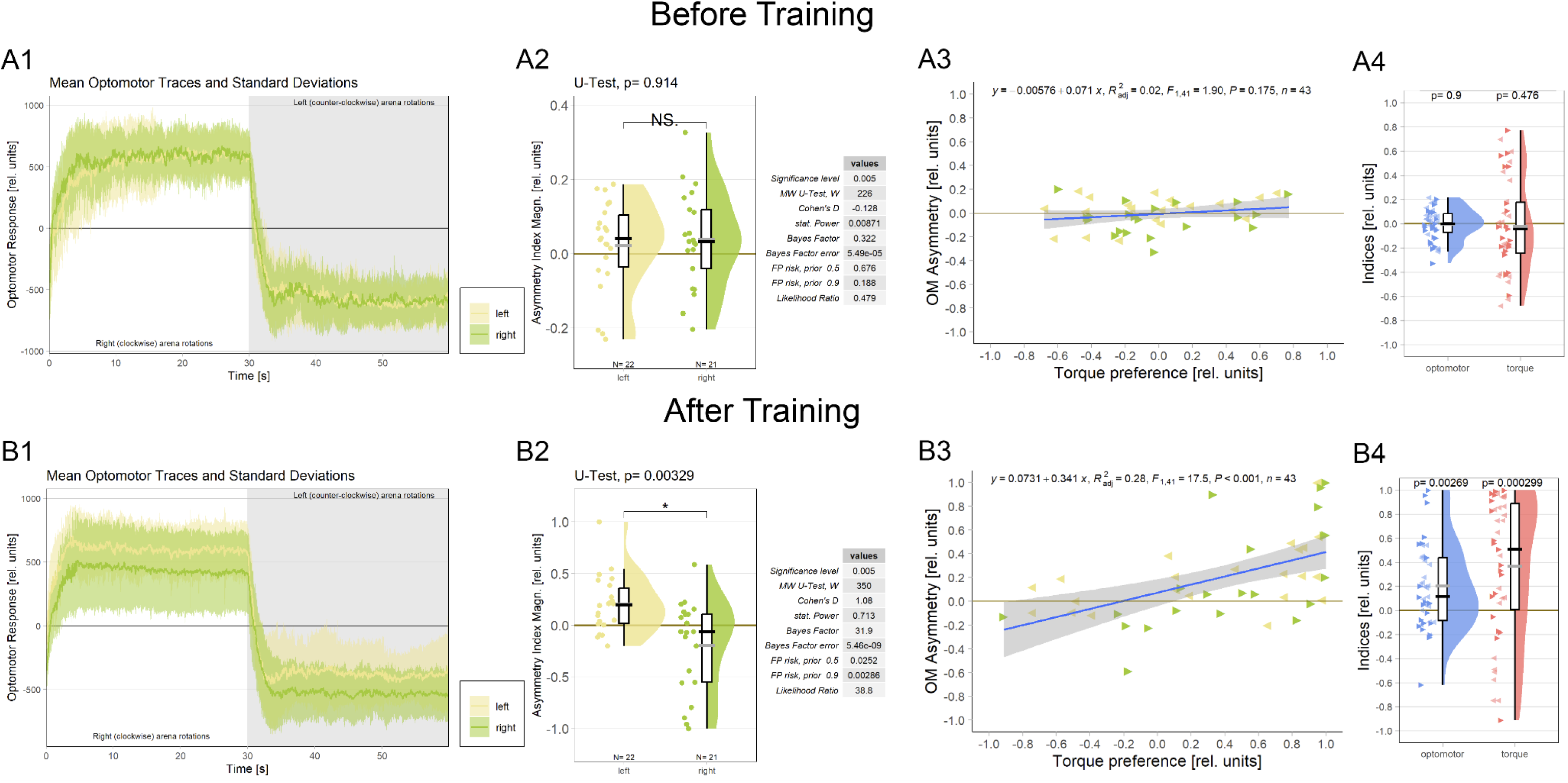
Associating one torque domain with heat changes optomotor behavior. A. Measurements before training **A1** Averaged optomotor traces of flies punished either on the ‘left’(yellow) or on the ‘right’ (green) torque domain. Both groups show similar response magnitudes in either direction of the optomotor stimulus. Errors are standard deviations. **A2** Optomotor asymmetry indices for flies punished either on the ‘left’(yellow) or on the ‘right’ (green) torque domain. The values for both groups spread around zero. Positive values indicate a shift towards positive (right-turning) torque. Both frequentist and Bayesian analyses are displayed to the right of the plots and indicate no difference between the groups. **A3** Regression analysis between torque preference and optomotor asymmetry. Optomotor values were adjusted such that positive values indicate a shift towards the unpunished torque domain. No significant correlation was observed. Left-pointing arrowheads (yellow) denote flies punished on left-turning torque and right-pointing arrowheads (green) denote flies punished on right-turning torque. **A4** Comparison of optomotor asymmetry (left, blue) and torque preference(right, red) indices. Here again, optomotor values were adjusted such that positive values indicate a shift towards the unpunished torque domain. Both measures vary around the zero point and Wilcoxon tests against zero are not significant (p-values above each plot). Left-and right-pointing arrowheads denote punishment directions as before. B. Measurements after training **B1** Averaged optomotor traces of flies punished either on the ‘left’ (yellow) or on the ‘right’ (green) torque domain. A reduction in the OMR magnitude can be observed on the punished, but not on the unpunished side. Errors are standard deviations. **B2** Optomotor asymmetry indices for flies punished either on the ‘left’ (yellow) or on the ‘right’ (green) torque domain. Positive values indicate a shift towards positive (right-turning) torque. The values for each group have now shifted towards the unpunished side compared to the values before training. Both frequentist and Bayesian analyses are displayed to the right of the plots and indicate a significant difference between groups. **B3** Regression analysis between torque preference and optomotor asymmetry. Optomotor values were adjusted such that positive values indicate a shift towards the unpunished torque domain. A significantly positive correlation was observed, such that higher torque preferences entailed larger optomotor asymmetry. Left-pointing arrowheads (yellow) denote flies punished on left-turning torque and right-pointing arrowheads (green) denote flies punished on right-turning torque. **B4** Comparison of optomotor asymmetry (left, blue) and torque preference (right, red) indices. Here again, optomotor values were adjusted such that positive values indicate a shift towards the unpunished torque domain. Both measures are shifted towards more positive values and Wilcoxon tests against zero are now significant for both variables (p-values above each plot). Left-and rightpointing arrowheads denote punishment directions as before. Data available at: 10.5283/epub.54804 (86)

Each direct steering muscle is innervated by one single identified MN (81). Thus, identifying the muscles reveals the identity of the corresponding MNs. To test for the expression of aPKC and *FoxP* in MNs that innervate direct wing muscles, we expressed UAS-GFP under the control of aPKC-Gal4 (Figure 3, second column) and RFP under the control of FoxP-lexA (Figure 3, third column). The signal of each fluorescent reporter was further enhanced by immunocytochemistry, and the active zone marker *bruchpilot* (*brp*; (66) was used to label the neuromuscular synapses in flight MN axon terminals (Figure 3, fourth column). In total, *Drosophila* is equipped with 12 flight steering muscles each of which likely contributes to a distinct function in flight control (82). The activation patterns of four of these muscles have been analyzed in response to optomotor stimulation during flight (first and second basalar muscles b1 and b2, as well as first pterale I muscle (i1) and third pterale II muscle (iii1) (82–84). A schematic of these 4 direct flight muscles plus five adjacent ones that were analyzed in this study illustrates their spatial arrangement, shapes, and depicts which ones are innervated by steering MNs that express both aPKC (green) and FoxP (magenta), only one of both, or neither of them (grey; Figure 3A1).

Representative maximum intensities projection views at 20x magnification allow us to visualize all nine flight steering muscles investigated in one field of view (Figs. 3A2-A4). Except the pleurosternal muscle 1 (ps1) and the posterior notal wing process muscles (hg), all other steering muscles are innervated by aPKC-positive MNs (Figs. 3A1, A2, green). In contrast, six of the nine steering muscles are innervated by *FoxP*-positive MNs, including the basalars b1 and b3, the pterale i1, iii3, and iii4, as well as the pleurosternal muscle ps1 (Figs. 3A1, A3, magenta). A subset of five flight steering muscles is innervated by MNs that express both genes required for operant self-learning. These are b1, b3, i1, iii3, and iii4 (Figure 3A).

Selective enlargements at 40x magnification were used to better visualize the axon terminals on those steering muscles of particular interest, with figure 3 panels B1-B4 focussing on the three muscles that insert at the basalar sclerite, and are thus named basalar 1-3 (b1-b3). Prominent labels of both aPKC (green) and *FoxP* (magenta) are present in the MNs to steering muscles b1 and b3, but the motor axon innervating b2 is devoid of *FoxP* signals (Figure 3, B1-B4).

Muscle b1 has been reported to fire once every wingbeat (83) during straight flight, during the transition from up-to downstroke. Optomotor stimulation causes phase shifts of b1 firing in the wing beat cycle, which in turn cause changes in wing beat amplitude both in the fruit fly *Drosophila* (83) as well as in the blowfly, *Calliphora* (85). The activity of b3 during flight has not been recorded electrophysiologically, but the muscle exhibits very similar morphological properties compared to b1. Both b1 and b3 are orientated similarly relative to the body axis and the wing hinge, and both are innervated by MNs with particularly large diameter axons (Figs. 3B1-B3) and particularly large active zones (Figure 3B4), suggesting similar functional roles. Steering muscle b2 is also innervated by large diameter axons (Figure 3C1) with large presynaptic active zones (Figure 3C4). Although some aPKC reporter label is detected in the MN to b2 (Figs. 3C1, C2, C4), labeling intensity is considerably fainter than that in b1 and b3 MNs (Figs. 3B1-B3). Fainter reporter labeling indicates weaker aPKC-Gal4 expression. The b2 MN is devoid of the *FoxP* reporter label (Figure 3C3). Although b2 has been reported to respond to optomotor stimulation, its phasic bursting responses correlate with rapid changes in wing beat amplitude as observed during body saccades (83).

Figure 3 panels C1-C4 move the field of view posteriorly and show the pterale 1 muscle i1 and the pterale 2 muscle iii1. Muscle i1 is innervated by a MN with faint aPKC (Figure 3C2) and faint *FoxP* (Figure 3C3) label. i1 has been reported to respond to optomotor stimulation, but its specific role in optomotor control remains largely unknown (83). Steering muscle iii1 shows some faint aPKC signal (Figure 3C2) but is devoid of any *FoxP* label (Figure 3C3) and does not participate in optomotor flight control (83).

The roles in flight control of the remaining four steering muscles (ps1, iii3, iii4, and hg; Figs. 3D1-D4) in *Drosophila* are not fully understood. However, iii3 and iii4 are innervated by MNs with aPKC (Figure 3D2) and *FoxP* (Figure 3D3) expression. Axon terminals on ps1 are *FoxP* positive (Figure 3D3) but lack any aPKC label (Figure 3D2). Active zones on steering muscle hg are visible through the *brp* label (Figure 3D4) but the motor axon on hg is devoid of both aPKC (Figure 3D2) and *FoxP* (Figure 3D3).

### 3.3. Self-learning breaks optomotor symmetry

Because it is not known at which torque meter reading the fly actually generates zero angular momentum, before each torque learning experiment, it is crucial to set the average of the asymptotic left and right optomotor response (OMR) magnitudes to zero (Figure 4A1, see Methods for details). This is to avoid an initial bias in the torque preference before training and with the assumption that the mid-point between the maximal OMRs roughly corresponds to flying straight (i.e, zero angular momentum). During the ensuing conditioning procedure, flies show spontaneous torque fluctuations that can reach or even exceed those elicited by optomotor stimulation in magnitude. Within each torque domain (i.e., ‘left’ or ‘right’, respectively), the temporal patterns (i.e., slow or fast) of torque fluctuations and their relative direction (i.e., more or less torque) are irrelevant for the heat stimulus as long as the zero point is not crossed: heat remains either on or off until the fly switches torque domains. Until this research, there was no reason to assume any relation between elicited OMRs and spontaneous torque fluctuations, neither conceptually nor anatomically. Now, however, the described role of the steering MNs co-expressing aPKC and FoxP (Figure 3) in large torque fluctuations elicited by optomotor stimuli prompted the hypothesis that there may be a neuroanatomical connection between spontaneous torque fluctuations and elicited OMRs after all: the motor neurons that innervate the steering muscles may be involved in both elicited and spontaneous torque fluctuations. If these neurons were indeed common to elicited OMRs and spontaneous torque fluctuations, we should observe a change in the OMRs after operant yaw torque learning.

To test this hypothesis, we analyzed the OMRs after training of a cohort of wild type Berlin flies (Figure 4B; control flies from a separate research project). We found that the asymptotic magnitude of the OMR on the punished side was reduced after operant training, while the OMR on the unpunished side remained unaltered compared to before training (Figure 4B1). Quantifying this observation revealed a significant difference in optomotor asymmetry after training (Figure 4B2) but not before training (Figure 4A2) between the two experimental groups.

An important question regarding the functional significance of this change in optomotor asymmetry is whether the amount of torque preference after training reflects the amount of optomotor asymmetry. There was no significant correlation between torque preference and optomotor asymmetry before training (Figure 4A3; as expected as the OMR was adjusted to be as symmetrical as practically possible). Once the flies have completed the training phase, torque preference becomes a significant predictor of optomotor asymmetry (Figure 4B3). These results corroborate our hypothesis that the wing MNs identified above, specifically those that have been shown to be involved in generating OMRs (82,83), constitute a site of the plasticity mediating operant self-learning in *Drosophila*. We also tested whether it would be sufficient to test for optomotor asymmetry after training as a proxy measure for torque preference. Before training, as expected, both measures showed similar values ranging around the zero point (Figure 4A4). After training, both measures did deviate from zero towards the unpunished direction, however, the effect was noticeably larger for torque preference than for optomotor asymmetry (Figure 4B4). Together with the correlation explaining about a third of the variance (Figure 4B3), these data may indicate that the MNs are an important, but not the only site of plasticity in this form of learning.

### 3.4. Spatial genome editing: *FoxP* likely not required in the brain

Optomotor analysis (Figure 4) suggests there may be additional sites of plasticity besides wing steering MNs. *FoxP*-positive neurons in the brain are straightforward potential candidates for such additional sites. Therefore, we performed spatial CRISPR/Cas9-based genome editing without restricting the manipulation to the adult stage. At the time of these experiments, no driver lines with *FoxP*-overlapping expression in the Saddle and Vest regions were available. Therefore, we tested driver lines expressing in the protocerebral bridge (PCB, Figure 5A), the PCB and adjacent central complex neuropils (see Materials and Methods for details, Figure 5B) and the dorsal cluster neurons (Figure 5C). Overlap of Gal4 expression patterns with FoxP expression was verified using the FoxP-LexA line (74,87) and FoxP-knockout efficiency was quantified previously (74). No self-learning impairment was observed following *FoxP* knockout in these brain areas, suggesting that *FoxP* expression is not necessary in these areas for operant self-learning. The results from these experiments, performed before aPKC was discovered as the likely gene necessary for operant self-learning, appear to corroborate the results of the aPKC/FoxP co-expression analysis in the brain: if both aPKC and FoxP are required for operant self-learning and there are no neurons in the brain that co-express both aPKC and FoxP, then knocking out either of them in the brain should not have any effects on operant selflearning.

**Figure 5:**
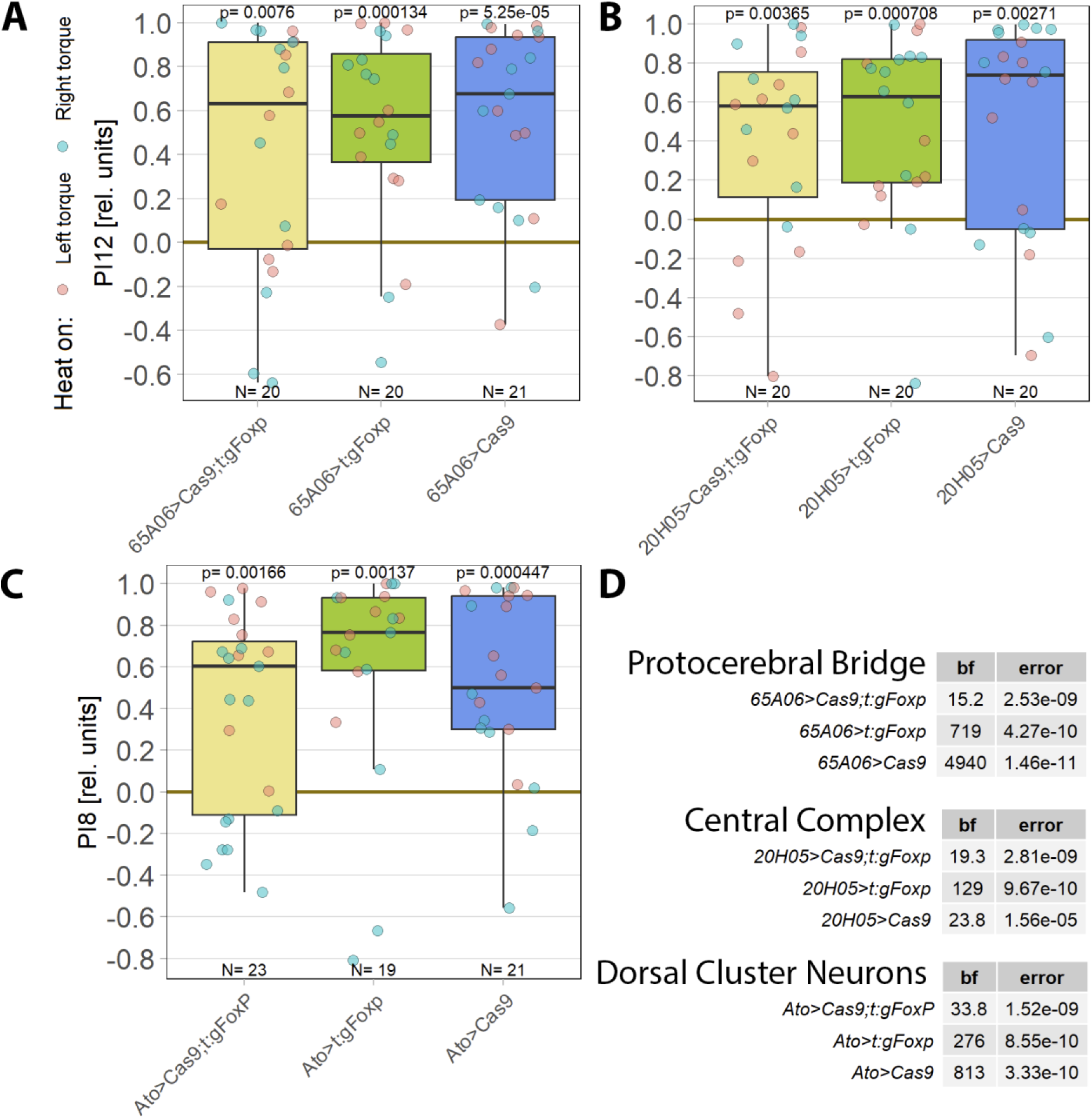
For operant self-learning, *FoxP* is likely not required in the brain. Performance indices for the first period after training (PI8/12) are plotted. Each colored dot represents a single fly experiment. Red dots denote experiments where the fly was punished on its left turning torque domain, blue dots denote flies that were punished on their right turning domain. Box plots denote medians, quartiles and non-outlier range. Statistical analyses test for differences of PIs against zero (above plots and D). A. Flies with *FoxP* knocked out in the protocerebral bridge (left, yellow) as well as the gRNA (middle, green) and Cas9 (right, blue) control flies showed high PIs, large Bayes factors (D) and small p-values, indicating their self-learning was intact. Data: 10.5283/epub.52956 (88) B. Flies with *FoxP* knocked out in the protocerebral bridge and additional components of the central complex (left. yellow) as well as the gRNA (middle, green) and Cas9 (right, blue) control flies showed high PIs, large Bayes factors (D) and small p-values, indicating their self-learning was intact. Data: 10.5283/epub.52951 (89) C. Flies with *FoxP* knocked out in the dorsal cluster neurons (left, yellow) as well as the gRNA (middle, green) and Cas9 (right, blue) control flies showed high PIs, large Bayes factors (D) and small p-values, indicating their self-learning was intact. Data: 10.5283/epub.52946 (89,90) D. Bayesian statistics for the three datasets.

We discovered that aPKC is required in MNs for operant self-learning (see above). As *FoxP* is also expressed in MNs, we knocked *FoxP* out in MNs using two different driver lines, C380-Gal4 and D42-Gal4. However, CRISPR/Cas9-mediated *FoxP* knockout in MNs disrupted flight-performance of manipulated flies to an extent that precluded any torque learning experiments. Our results above (Figure 3) will now allow us to select more specific driver lines, expressing only in the identified wing steering MNs (91), which may yield *FoxP* knock-out flies (or any other manipulation) with sufficient flight performance.

### 3.5. Temporal genome editing: Self-learning impaired only after a 2-week *FoxP* knockout

*FoxP* was shown to be important for normal development (74,92). Strong motor impairments have been reported after a developmental *FoxP* knockout, for instance rendering the animals unable to fly (see above). Since the ability to fly is a basic requirement for torque learning, we used a CRISPR/Cas9-based approach to pan-neuronally knock out *FoxP* in adult flies. No effect was observed on self-learning two days after the start of the knockout induction (Figure 6 A), but waiting for 14 days after the pan-neural knock-out yielded a significant self-learning impairment (Figure 6B). As a test seven days after induction was also without effect (DOI: 10.5283/epub.52965), self-learning remains functional for at least 7-14 days after cessation of *FoxP* gene transcription in all neurons.

**Figure 6:**
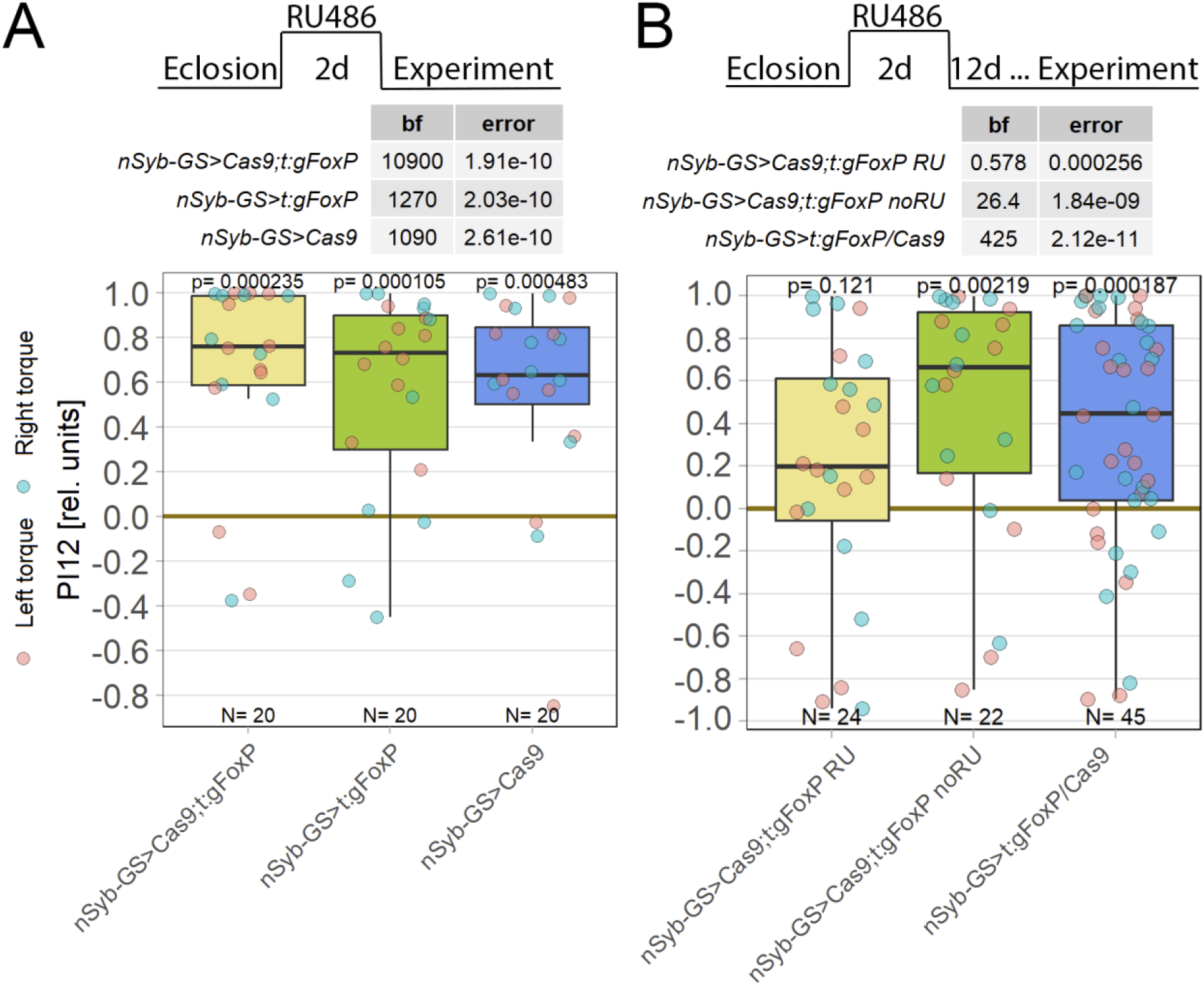
Knockout of *FoxP* in adult *Drosophila* shows learning impairments only after 14 days. Plotted are performance indices for the first period after training (PI12). Each colored dot represents a single fly experiment. Red dots denote experiments where the fly was punished on its left turning torque domain, blue dots denote flies that were punished on their right turning domain. Box plots denote medians, quartiles and non-outlier range. Statistical analyses test for differences of PIs against zero. A. Self-learning two days after *FoxP* knockout induction with RU486. Experimental animals (left, yellow) as well as gRNA (middle, green) and Cas9 (left, blue) control animals showed all high PIs as well as large Bayes Factors above 1000 and p-values below 0.005, indicating that all groups showed unimpaired self-learning. Data: 10.5283/epub.52963 (93) B. Self-learning 14 days after *FoxP* knockout induction with RU486 and 12 days after cessation of RU486 administration. Experimental animals (left, yellow) showed low PIs as well as a Bayes Factor of less than one together with a large p-value, whereas both the genetic control animals without RU486 treatment (middle, green) and the pooled RU486-treated gRNA and Cas9 controls (right, blue) showed high PIs, large Bayes Factors and p-values smaller than 0.005, indicating an impairment in self-learning only in the experimental group. Data: 10.5283/epub.52964 (94)

### 3.6. *aPKC* acts via non-canonical pathways in self-learning

Having established that *aPKC* is required in *FoxP*-positive MNs, we sought to identify further components of the *aPKC*-dependent plasticity underlying operant self-learning in *Drosophila*. It is not uncommon for plasticity mechanisms in the adult animal to recruit genes with a function during neuronal development (95–97). With PKCs being notorious for being able to compensate for genetic manipulations (69–73), one reliable countermeasure has proven to shorten the time period between manipulation and testing sufficiently to ensure compensation has no time to take place (21,31). Following this tried-and-tested approach, our manipulations of prominent PKC interaction partners were thus restricted to adult neurons. One such prominent interaction partner of *aPKC* during *Drosophila* nervous system development is *bazooka* (*baz*), a crucial component of the highly conserved PAR complex (98,99). However, knocking out *baz* in all adult neurons did not disrupt operant self-learning (Figure 7A), suggesting that the Par complex signaling pathway is not involved in operant self-learning. A second prominent aPKC interaction partner is the kidney and brain protein (KIBRA), which acts in the conserved Hippo pathway (100,101) and also proposed to be involved in learning/memory (102–106). Knocking out KI-BRA in all adult neurons also did not disrupt operant self-learning (Figure 7B).

**Figure 7:**
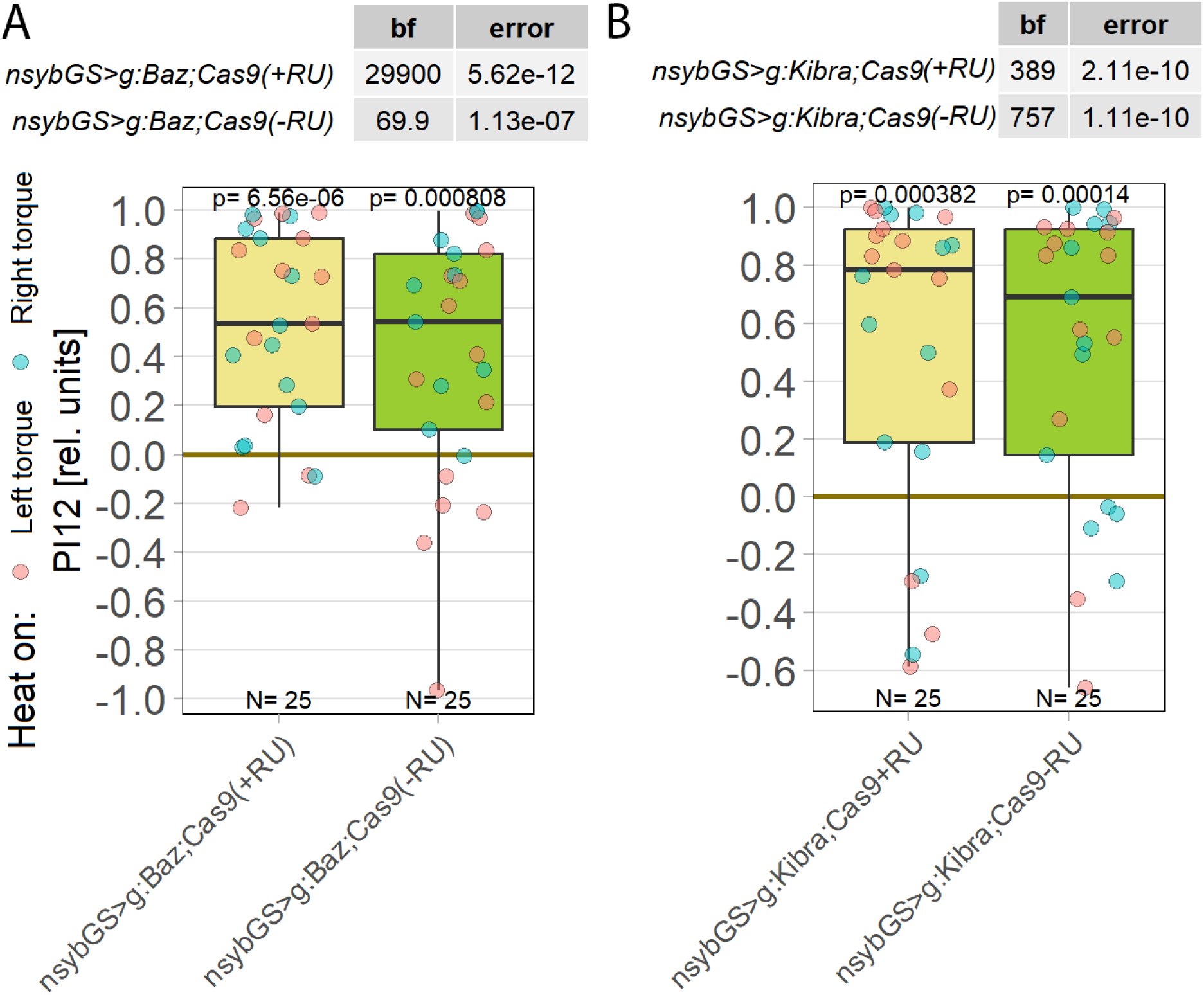
Neither *bazooka* nor KIBRA are involved in operant self-learning. Plotted are performance indices for the first period after training (PI12). Each colored dot represents a single fly experiment. Red dots denote experiments where the fly was punished on its left turning torque domain, blue dots denote flies that were punished on their right turning domain. Box plots denote medians, quartiles and non-outlier range. Statistical analyses test for differences of PIs against zero. A. Knocking out *bazooka* using CRISPR/Cas9-mediated genome editing in adult animals had no effect on operant self-learning. Both flies that were fed the steroid hormone RU486 (left, yellow) and the genetically identical flies without the hormone (right, green), showed high PIs, low p-values and high Bayes Factors, indicating a preference for the unpunished torque domain. Data: 10.5283/epub.52947 (107) B. Knocking out KIBRA using CRISPR/Cas9-mediated genome editing in adult animals had no effect on operant self-learning. Both flies that were fed the steroid hormone RU486 (left, yellow) and the genetically identical flies without the hormone (right, green), showed high PIs, low p-values and high Bayes Factors, indicating a preference for the unpunished torque domain. Data: 10.5283/epub.53685 (108)

## 4. Discussion

Operant self-learning in *Drosophila* is a form of motor learning that appears to be conserved among bilaterians. The transcription factor *FoxP* is involved in various forms of motor learning in chordates (5–9,11,13,16–18), as is PKC (22–24,109–112). PKC is also involved in motor learning in the feeding behavior of the lophotrochozoan *Aplysia* (*113*) and both are involved in motor learning in the ecdysozoan *Drosophila* (this work and (20,21,31). This wealth of evidence supports what has been called ‘deep homology’ for motor learning in bilaterians (12).

In this work, we present new insights into the neurobiological mechanisms underlying this form of motor learning.

### 4.1. Self-learning requires *aPKC* in *FoxP* wing steering motor neurons

While previous evidence suggested that some PKC activity was required in some MNs for operant self-learning (21), it was not clear which PKC gene was involved and in which MNs. We now hypothesize that *aPKC* activity is required in aPKC/*FoxP*-positive direct wing steering MNs in the ventral nerve cord (VNC). Our analysis of these direct wing muscles, responsible for, e.g. wing beat amplitude (a major contributing factor to yaw torque) (114), revealed *aPKC/FoxP* co-expressing MNs innervating a specific subset of these muscles (Figure 3), corroborating our hypothesis.

### 4.2. Motor neurons involved in generating yaw torque co-express aPKC and *FoxP*

Specifically, the basalar muscle b1 which is known to regulate wing beat amplitude during optomotor stimulation through phase shifts of its action potential within the wing beat cycle (83) is innervated by a MN with very strong aPKC and *FoxP* expression in all 11 animals analyzed. Similarly, b3 is also innervated by a MN with very strong aPKC and *FoxP* expression and the b3 muscle is reported to increase activity when the ipsilateral wing decreases its amplitude (115). The b1 and the b3 muscles thus both share similar sizes and morphologies, are innervated by MNs with particularly thick processes and presynaptic active zones, insert at opposite sides of the basalar sclerite and act in a push-pull fashion in controlling wing beat amplitude (115), a major contributor to yaw torque. A recent comprehensive re-analysis of all steering muscles has confirmed the antagonist role b1 and b3 play in generating yaw torque (116). Although the MN innervating the third basalar muscle, b2, also expresses aPKC along the axon, we did not detect aPKC signal in axon terminals or active zones, and it does not show *FoxP* label, thus ruling out a function of b2 in aPKC/FoxP mediated operant self-learning. In contrast to b1 and b3, b2 firing is not linked to the wingbeat cycle; instead, it fires in bursts during turning maneuvers (82). Corroborating the conclusion that it is not involved in mediating operant self-learning, b2 is silent during straight flight, but it is likely involved in regulating body saccades (Heide, Götz, 1996), which are by themselves not relevant for controlling the heat in our experiments. For the basalar muscles the emerging picture is that b1 and b3 are the top candidates for self-learning because they likely regulate wingbeat amplitude and thus also yaw torque on a wingbeat cycle by cycle basis (116). Strikingly, the MNs innervating these two muscles show high levels of aPKC and FoxP expression. Although much less is known about the pterale muscles, and neither iii3 nor iii4 (both with aPKC and FoxP co-expression) have been recorded electrophysiologically during flight, a similar picture begins to emerge. The MN to the left muscle i1 is known to be active during right turns (83) and co-expresses aPKC and FoxP, whereas iii1 reportedly (83) does not participate in OMRs and is innervated by a MN with weak aPKC and no *FoxP* label. Also these roles of steering muscles have recently been confirmed (116). In summary, flight steering muscles that are known to regulate yaw torque in response to horizontal optomotor stimulation receive aPKC and *FoxP* positive motor innervation, while those not involved in yaw torque production are not innervated by such doubly labeled MNs. Because endogenous yaw torque fluctuations, such as those rewarded/punished in our operant experiments also have to use these MNs, it becomes clear that OMRs and operant yaw torque learning share these steering motor neurons as a common component. Future experiments with (likely soon to be available) more specific driver lines will be able to test the effect of aPKC/FoxP manipulations in specific subsets of steering MNs.

### 4.3. Operant self-learning modifies optomotor responses

From this realization, the question arises if there are other shared neural components between OMRs and operant learning in addition to the doubly labeled steering MNs. Evidence suggests that the steering commands for OMRs are communicated directly from visual areas in the brain via (both identified and yet to be identified) descending neurons with direct synaptic connections onto the steering MNs in the VNC (117,118). This suggests that in the ventral nerve cord, the steering motor neurons are the only neurons that OMRs and operant self-learning share. This evidence entails that any additional overlap would need to be localized in the brain, where no neurons expressing both aPKC and FoxP. seem to exist (Fig. 2). The remote hypothetical possibility remains that OMR plasticity after operant learning may be caused by non-aPKC/Foxp-dependent plasticity mechanisms in the same visual parts of the brain where also the steering commands for OMRs are computed. Future research will address this possibility. The currently available evidence thus points towards the steering MNs we have identified as the only set of neurons where a modification due to operant learning could have an effect on OMRs. The fact that we have observed precisely such an effect (Fig. 4), only leaves the doubly labeled steering neurons (Fig. 3) as the sites for the aPKC/FoxP mediated plasticity mechanisms underlying operant yaw torque learning.

Then again, only about a third of the variance in the torque preference after training can be explained by this optomotor asymmetry. This result explains the observations that there are flies with a strong conditioned torque preference but with an optomotor asymmetry in the opposite direction, as well as flies with a weak conditioned torque preference and large optomotor asymmetry (Figure 4B3). Clearly, plasticity in steering MNs appears to be important, but it does not reflect the entirety of the learning processes. On the other hand, it may be that the strong OM stimuli we use here overshadow potentially larger asymmetries in OMRs. We will test this in future experiments using weaker OM stimuli after operant self-learning.

Plasticity in steering movements such as OMRs have been observed before, such as in classic “inversion goggles” experiments where the coupling between the fly’s movements and the environment was reversed (119) or in a more recent experiment revealing adaptation processes (120). It has long been recognized that insects with asymmetrical wing damage need to adjust the neural commands for generating torque to compensate for the changed physical torque (e.g., (117,118,121–125)). Plasticity in wing steering motor neurons provides a potential mechanism for such adjustments.

### 4.4. Motoneuron plasticity mediates operant selflearning

Taken together, these results converge on the hypothesis that plasticity in MNs that innervate the direct muscles involved in generating yaw torque, but not other steering MNs, mediates an important aspect of operant self-learning in *Drosophila*. The importance of MN plasticity is emphasized by *FoxP*-dependent plasticity apparently not being required in the brain (Figure 5). This MN plasticity could either be implemented by (a) postsynaptic plasticity of the input synapses to these MNs, postsynaptic because the MNs but not the interneurons express the proteins required for self-learning, or (b) on the level of the intrinsic excitability of flight steering motoneurons, or (c) on the level of the output synapses to the respective steering muscles. However, the mechanism and subcellular localization of self-learning in MNs remains to be determined.

A second interesting, yet unstudied population of *aPKC/FoxP* co-expressing neurons resides in the abdominal neuromer of the VNC. As flies use their abdomen analogously to a rudder during turns in flight (126,127), involvement of these neurons seems plausible in addition to wing MNs.

While few studies in insects have shown MN plasticity, the *Aplysia* sensorimotor synapse is a classical model for research on plasticity mechanisms. There is a rich literature on MN plasticity in this preparation, some of which reports PKC-dependent mechanisms (104,128–138). Also in mammals (including humans) MN plasticity in the spinal cord is a readily observable phenomenon in nonclinical and clinical settings (139–148). The discovery of aPKC-dependent plasticity in *Drosophila* MNs expands this body of literature to a genetically tractable organism and inasmuch as clinical practice relies on MN plasticity, may even help instruct the development of clinical applications.

Until our work, PKC activity had only been shown to be important for memory consolidation/maintenance in world-learning experiments in flies, but not for learning/acquisition (149,150). Also in *Drosophila* at the torque meter, PKCs are dispensable for world-learning (31). The literature on PKCs in learning and memory in other animals is complex and multifaceted. In some preparations, PKC isoforms are required during memory maintenance, in some also during acquisition and in others different isoforms distinguish between acquisition and consolidation (69,70,102,104,129,134,136,151–159). As the manipulations in our experiments lasted throughout training together with the tests immediately following training and we did not test for long-term memory, we can only ascertain that aPKC is required in a very narrow time window of minutes around training. Future research will address whether aPKC must be present during training, test, or both.

Notably, there is one other preparation where PKC activity is involved and which is also conceptually most closely resembling the one we used here, operant reward learning in *Aplysia* feeding behavior. However, in this preparation, the calcium-dependent Apl-I PKC not the atypical Apl-III PKC appears to be mediating the plasticity (113). Given the degeneracy between the different PKC genes and the fact that they can not only compensate for long-term PKCi-mediated inhibition (this work and (21), but also for each other (69), more research is needed to elucidate how these different mechanisms of plasticity evolved and are related to each other.

### 4.5. Self-learning may be mediated by a non-canonical *aPKC/FoxP* pathway

The observation that neither *bazooka* (*baz*) nor the kidney and brain gene (KIBRA), two prominent interaction partners of aPKC (98–104,106,160,161), showed an effect on operant self-learning when they were knocked out in the adult nervous system (Figure 7), raises questions about the effectiveness of the CRISPR/Cas9 method in these cases. In particular, one may question the approach of temporally limiting the manipulation to adult neurons. Although this had been effective with previous PKC manipulations using an inhibitory peptide, PKCi (21,31), it only yielded an intermediate effect with a CRISPR-mediated knockout targeting aPKC (Figure 1B). In contrast, temporally uncontrolled knockout of aPKC in MNs of the VNC (Figure 1C) proved surprisingly effective, given expectations from prior experience.

Nonetheless, we are confident that both manipulations successfully knocked out each of these genes in a way sufficient to induce a phenotype, if either were necessary for self-learning. First, we found that the *baz* knockout, rather than impairing self-learning, actually *increases* learning performance (manuscript in preparation, thesis: (87)), suggesting the PAR complex may be sequestering *aPKC* and thereby limiting its availability for self-learning. Thus, these data suggest that *baz* is indeed not directly involved in mediating the aPKC activity contributing to plasticity in steering MNs; instead, it may be binding aPKC in the PAR complex, preventing it from playing its role in MN plasticity (162). Second, our KIBRA knock-out had a severe effect on flight performance when elicited during development, suggesting that also this manipulation was, in principle, effective. That being said, without clear evidence that the *baz* and KIBRA proteins are completely absent, these results remain suggestive rather than conclusive. In addition to protein-level analysis of CRISPR efficacy (as we have performed in CRISPR-mediated FoxP knock-out (74)), future experiments will use FoxP-iB and c380/D42 drivers to drive baz and KIBRA knockouts. The forthcoming full description of the KI-BRA gRNA line by the Krahn laboratory will also help bolster or refute the results we have obtained.

Interestingly, at least during development, *baz* and KIBRA have been reported to have opposite effects on the function of *aPKC*, with the PAR complex (*baz*) and the Hippo pathway (KIBRA) mutually inhibiting each other (163). While *baz* is thought to mediate *aPKC* activity, KIBRA is thought to exert negative regulatory effects on *aPKC*. Thus, knocking out each one of them should have revealed a decrement in self-learning in at least one of them, if the processes during development were recapitulated during self-learning. On the other hand, experiments in which KIBRA has been shown to be involved in learning/memory have suggested a positive rather than a negative regulatory role (104,154), albeit with an emphasis on long-term memory rather than learning (164). Whichever way *aPKC* may be interacting with components of these canonical pathways, the literature predicts that at least one of our manipulations should have revealed a decrement in operant self-learning. The fact that this prediction was falsified may suggest that *aPKC* exerts its function in a non-canonical manner in operant self-learning plasticity. We are currently pursuing research into the possibility that *bazooka* may be a negative regulator of aPKC activity (162) during operant self-learning.

### 4.6. Persistent *FoxP* effect on operant self-learning in adults

As *FoxP* mutants are impaired in operant self-learning (20), two hypotheses about the role of this prominent transcription factor arise. First, *FoxP* may be directly involved in the learning process via some unknown, cytosolic function. A transcription factor function appears unlikely, because of the short duration of our experiments. Second, *FoxP* may exert its effects as a developmental regulator, being crucial for the development of the circuits mediating operant self-learning. As the developmental role of FoxP genes is well documented (7,14,74,92,165,166) and there are few domains in the gene that would lend themselves to a hypothetical cytosolic function, the latter hypothesis appeared more plausible. The result that adult knockout of *FoxP* had no immediate effect on operant self-learning (Figure 6A) seemed to corroborate this hypothesis. In contrast, supporting a continued role of FoxP genes in motor learning also after development are data from songbirds where *FoxP2* gene expression is not only regulated by singing (15,167–169), but where normal *FoxP2* expression is necessary in adults to maintain learned song (8,170). To also test the second hypothesis, we aged the flies after the *FoxP* knockout and tested them 7 and 14-days later. Our results (Fig. 6) suggest a role for adult *FoxP* expression in maintaining operant self-learning capabilities after development, analogous to the role of FoxP2 in songbird vocal learning (8,170). There are three possible explanations for this result: For one, the genes regulated by *FoxP* (*171*) may continue to exert their functions for this amount of time also without the FoxP protein present. Second, the half-life of the FoxP protein may be long enough. Third, a combination of these two explanations. Further research is needed to distinguish between these options.

## Data availability

### Underlying data, figures

1A: 10.5283/epub.52958

1B: 10.5283/epub.52957

1C: 10.5283/epub.52944

2: 10.5281/zenodo.10047941

3: 10.5281/zenodo.10606166

4: 10.5283/epub.54804

5A: 10.5283/epub.52956

5B: 10.5283/epub.52951

5C: 10.5283/epub.52946

6A: 10.5283/epub.52963

6B: 10.5283/epub.52964

7A: 10.5283/epub.52947

7B: 10.5283/epub.53685

License: CC0

## Software availability

Latest source code available from: https://github.com/brembslab/DTSevaluations

Software controlling experiments and collecting data: https://www.doi.org/10.5281/zenodo.7102195

Software evaluating data: https://www.doi.org/10.5281/zenodo.10041052 Data model:

10.5281/zenodo.10041052

Python script keeping local and public datasets synchronized:

10.5281/zenodo.7101741

License: GPL-3.0

## Acknowledgements

We thank Thomas Kopp in the electronics workshop of the University of Regensburg, without whose tireless support this work could not have been accomplished. We are indebted to Michael Krahn for very helpful feedback on PKC interaction partners and for sending us his KIBRA gRNA flies before publication. We are grateful to several anonymous peer-reviewers as well as the named reviewers of this submission (Efthimios M C Skoulakis, Katrin Vogt and Wayne Sossin) who have helped improve the manuscript substantially. We thank the DFG for financial support: DFG BR 1892/17-1. This work is dedicated to the late Jochen Pflüger, at whose memorial CD and BB agreed to collaborate on motor neurons.

